# The nuclear oncoprotein SET is necessary for MLL/KMT2A binding and transcriptional elongation

**DOI:** 10.64898/2026.02.26.708410

**Authors:** Maria-Paz Garcia-Cuellar, Robert K. Slany

## Abstract

The nuclear oncoprotein SET (patient “SE” translocation) has been implicated in the etiology of MLL/KMT2A-fusion induced leukemia. Here we examine the details of this dependency in murine, primary hematopoietic cells. Experiments demonstrated *Set* as downstream target of HoxA9 and a direct interactor of Mll/Kmt2A. Mll/Kmt2A and Set globally co-bound promoter regions. Impairing Set expression induced a metabolic shift towards oxidative phosphorylation phenocopying a knockdown of Mll/Kmt2A fusion targets. Set acted predominantly as transcriptional activator driving a pro-proliferative gene expression program with features indicative for Mll/Kmt2A involvement. Molecularly, Set depletion caused dissociation of Mll/Kmt2A from chromatin accompanied by a selective loss of elongating RNA PolymeraseII Ser2-P. Concomitant with a function of Set as inhibitor of protein phosphatase 2A (PP2A), specific recruitment of PP2A to the Meis1 promoter, a known Mll/Kmt2A target, inhibited transcription in reporter assays and in a natural chromatin environment. We identified Mitogen and stress induced kinase 1 (Msk1) as potential substrate protected by Set from dephosphorylation. Active and phosphorylated Msk1-P colocalized with Mll and disappeared from chromatin upon Set depletion. Biochemically, Msk-1 bound directly to Mll/Kmt2A as well as to menin, a known Mll/Kmt2a tethering factor. Loss of Set/Mll/Msk1 selectively affected H3K14 acetylation at promoters and this could be partially attributed to the reduced presence of the histone acetyltransferase Moz/Kat6a. Finally, we show that kinase and menin inhibitors cooperate in leukemia cells indicating that the relay function of Mll/Kmt2A, allowing control of hematopoiesis by cellular signaling, is retained in MLL-fusion proteins.

## Introduction

The nuclear oncoprotein SET (acronym for patient SE translocation) is an evolutionary highly conserved, multifunctional protein that is frequently involved in malignant transformation. Despite confusion in literature, SET is not related to SET methyltransferase-domain containing (<u>S</u>uppressor of variegation, <u>E</u>nhancer of zeste, <u>T</u>rithorax) proteins. SET is a predominantly nuclear protein with an unusual structure and no known enzymatic activity (figure 1A). At the C-terminus, it carries an unstructured, highly acidic domain that may serve as an inert binding decoy. SET is detectable in cells in various isoforms originating from alternative transcript start sites and differential splicing. SET has been reported to play a role in a puzzling variety of physiological functions ranging from cancer to Alzheimer disease (for reviews see ^1, 2^). While the underlying mechanisms are far from clear, the major and best researched activity of SET is inhibition of the threonine-serine phosphatase PP2A (protein phosphatase 2A) ^3^.

**Figure 1:**
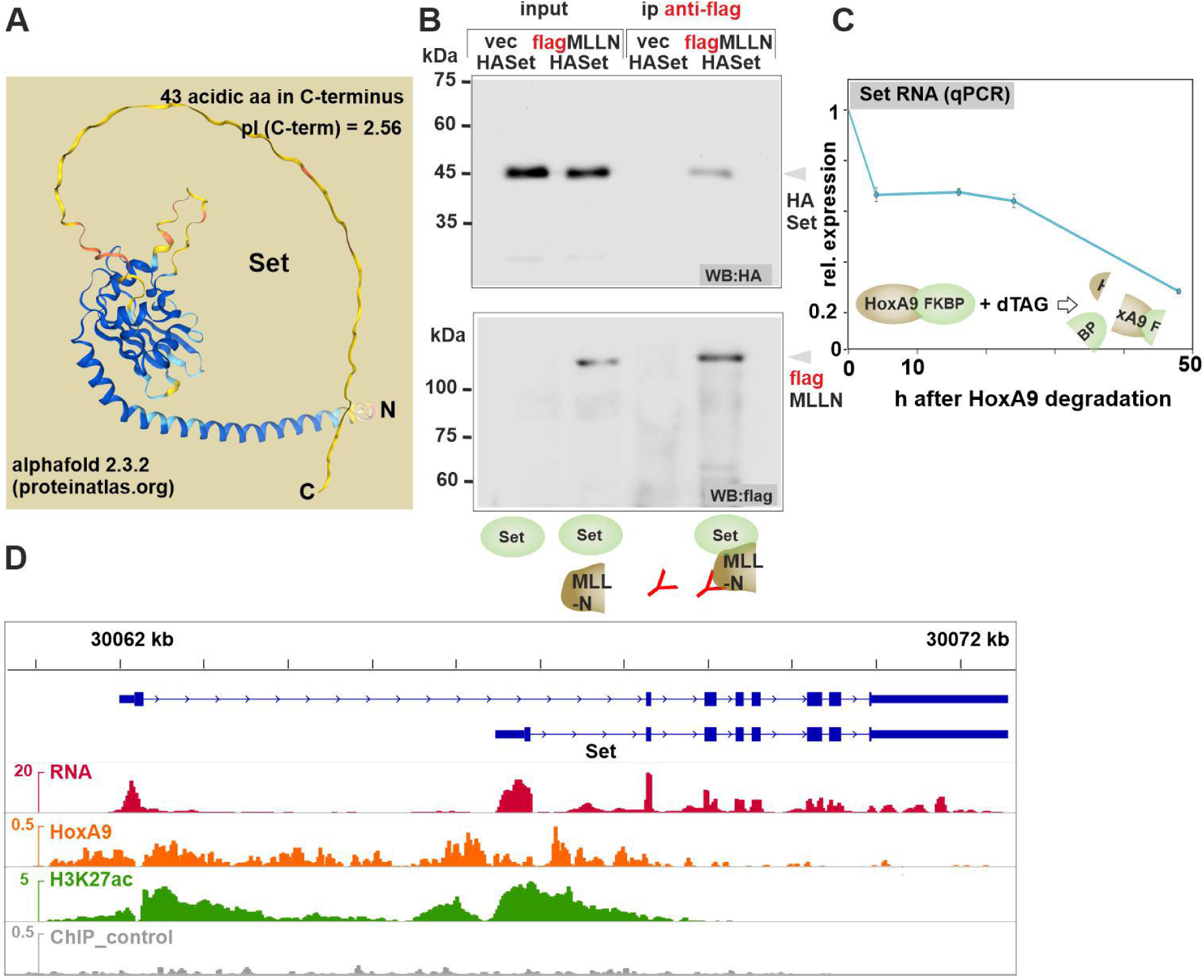
Set interacts with MLL-N and *Set* transcription is under control of HoxA9. A: Structure of Set according to alphafold 2.3.2, Source: www.proteinatlas.org B: Coimmunoprecipitation of Set and the N-terminus of MLL conserved in fusion proteins. Immunotagged versions of Set and MLL-N as indicated were coexpressed in 293T cells. IP was done with MLL-N as bait an precipitates analyzed by western blot. C: Set expression was quantified by q RT-PCR in cells immortalized by a degradable HoxA9-FKBP construct during a time course of HoxA9 depletion initiated by addition of dTAG13. D: IGV snapshot of RNA expression and HoxA9 binding/H3K27acetylation at the Set-locus in myeloid precursors.

Overexpression of SET in malignancies is frequently associated with a dismal prognosis and the SET-cancer connection is particularly obvious in acute leukemia with an emphasis on MLL/KMT2A associated disease (While technically deprecated, MLL still is the more frequently used designation and hence will be used throughout this article). Originally, SET was discovered as fusion partner of NUP214 (originally named CAN) in a case of myeloid leukemia ^4^. Later, SET was reported to bind to the methyltransferase MLL/KMT2A ^5^. SET transcription was found to be controlled by MLL fusions ^6^ and by HOXA9 ^7^. More evidence for an implication of SET in leukemogenesis comes from the investigation of SETBP1 (SET binding protein 1). SETBP1 binds to and protects SET from degradation ^8^. Overexpression of SETBP1 alone is sufficient to immortalize hematopoietic precursor cells by upregulation of HOXA9 and HOXA10 ^9^. In myelodysplastic syndromes, SETBP1 mutations are frequent and cause overexpression of MECOM, a repressor of myeloid differentiation ^10^. SETBP1 contains two AT-hook DNA-binding domains and colocalizes with MLL complexes to aberrantly expressed chromatin. Finally, two recent articles ^11, 12^ implicate SET in the function of MLL fusion proteins. A knockdown of SET impaired the transforming activity of MLL fusions by reducing target gene transcription. MLLr (MLL recombined) cells were also sensitive to administration of FTY720, a pleiotropic acting substance that, amongst other effects, disrupts the ability of SET to inhibit PP2A. While inhibition by FTY720 was mainly explained by a reduction of Myc phosphorylation, further molecular details of SET function in the context of MLL activity remained largely unexplored.

Here we investigate this question more in detail and we find that chromatin association of MLL and specific induction of transcriptional elongation is highly dependent on Set-dependent maintenance of phosphorylation. This indicates MLL functions as a relay that receives signals to control gene expression while Set acts as modifier of signal strength.

## Results

### Set binds to the N-terminus of MLL and it is a direct target of HoxA9

While previous publications implicated Set in MLL function, there is only scarce evidence for a direct interaction. Therefore, we first confirmed Set binding to the N-terminus of MLL retained in fusions (MLL-N) by co-immunoprecipitation (figure 1B). We also corroborated the regulation of *Set* by HoxA9 in HoxA9-FKBP degron transformed myeloid precursors. *Set* transcripts rapidly dropped by 40% four hours after addition of degradation inductor dTAG13 suggesting that at least part of *Set* transcription is under direct control of HoxA9 (figure 1C). This was supported by ChIP where HoxA9 protein prominently bound to H3K27-acetylated areas around the transcriptional start sites of both annotated *Set* transcripts (figure 1D).

### Genome wide colocalization of Set and Mll at promoter regions

Next, we wanted to determine the genome-wide localization of endogenous Set and MLL. Set lacks a DNA binding domain and probably does not contact DNA directly, making crosslinking with a “zero-length” crosslinker like formaldehyde difficult. Therefore, ChIP was performed with EGS (Ethylenglycol bis Succinimidyl-Succinat) that provides a 12-atom spacer. With this modified ChIP procedure, we identified 620 high confidence Set peaks (examples in figure 2A) that colocalized with Mll and H3K27acetylated chromatin. General congruence of Set and Mll presence was corroborated by metagene analysis (figure 2B, C) and occurred overwhelmingly in regions annotated as promoters (figure 2D).

**Figure 2:**
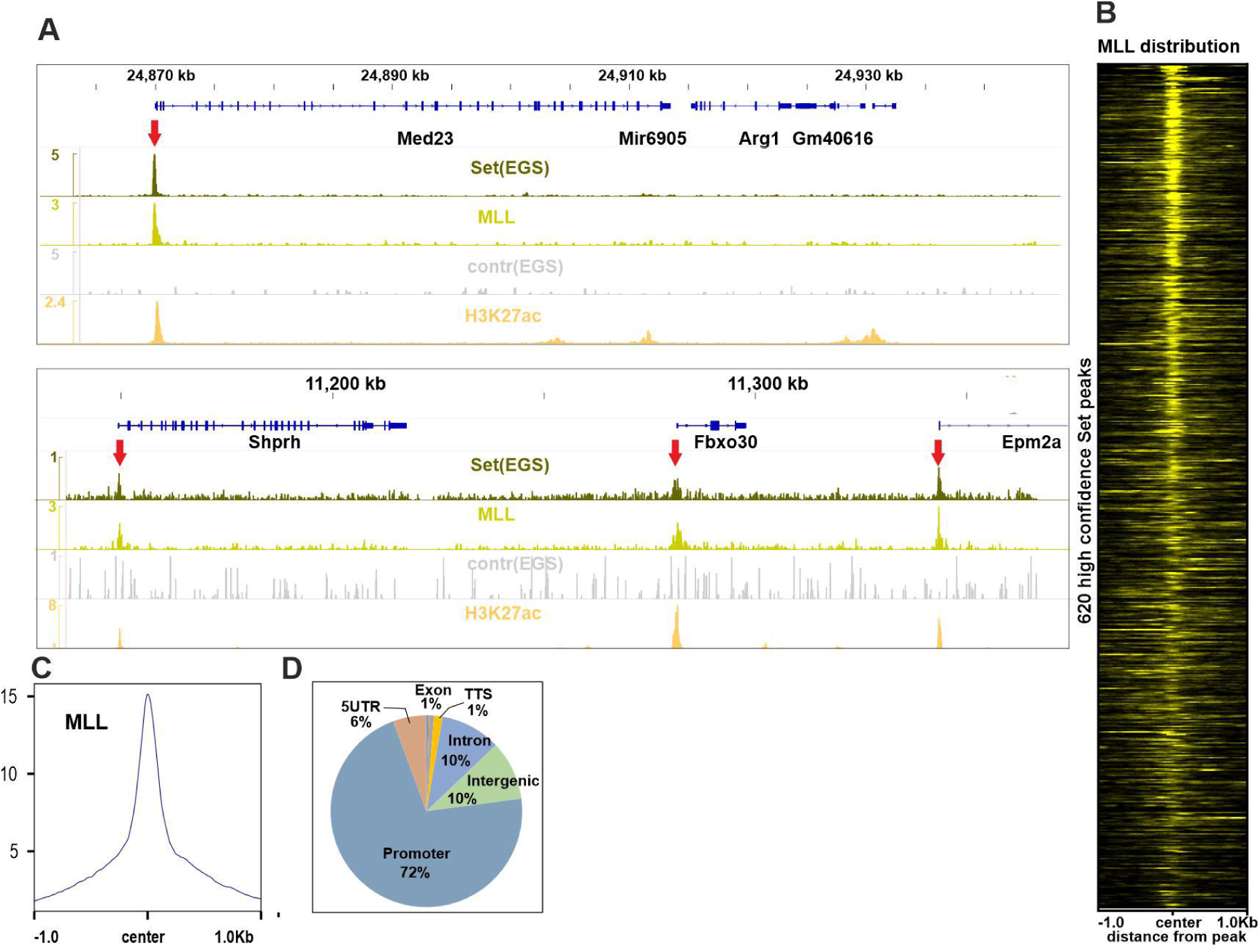
Set and Mll globally colocalize to promoter regions. A: Integrated genome viewer (IGV) snapshots of Set, Mll and H3K27ac profiles. Set ChIP was done after crosslinking with EGS with a specific input control included. B: Metagene plot showing Mll distribution around Set peaks. The plot incorporates 620 Set-peaks identified with high confidence (6-fold enrichment over input, 8-fold enrichment over local background). C: Summary plot of Mll distribution around Set binding sites D: Functional annotation of Set peaks.

### Set knockdown causes a metabolic phenotype

To investigate the consequences of a loss of Set activity in myeloid precursor cells, we applied an inducible shRNA knockdown system (figure 3A). Constructs delivered by lentiviral transduction contained a Set-specific (shSet) or a non-targeting (luciferase, pLEPIR) hairpin allowing to control for unspecific effects of tetracycline induction. As the shRNA hairpin is integrated into the 3’UTR of a truncated *LNGFR* gene (low affinity nerve growth factor receptor, CD276) shRNA positive cells could be identified by FACS and positively selected. The shRNA efficiently reduced all five different isoforms of Set as detected by Western (figure 3B, left panel). Cells with depleted Set were rapidly lost during culture (figure 3B, right panel). Therefore, experiments were conducted with magnetically selected shRNA-positive cells cultured for a maximum of three days. Unexpectedly, while standard MTT tests suggested a proliferation defect after Set knockdown, absolute cell numbers were unaffected (figure 3C) a phenomenon that can appear if cells differ in availability of NADH/H+ for MTT reduction. Phenotypically, depletion of Set strongly reduced medium acidification during culture (figure 3D). This could be traced to a reduction of glucose consumption and lactate production in Set knockdown cells compared to controls (figure 3E). A highly reminiscent phenotype was described before in MLL-ENL transformed cells depleted for the MLL-ENL target gene *Ptpb1*^13^. Cells lacking Ptbp1 (or its binding partner Hnrnpa1) change splicing of pyruvate kinase M (PKM) to an isoform that favors oxidative phosphorylation over glycolysis. Consequently, NADH/H+ is predominantly used for ATP synthesis rather than for lactate synthesis (schematic depiction in figure 3F).

**Figure 3:**
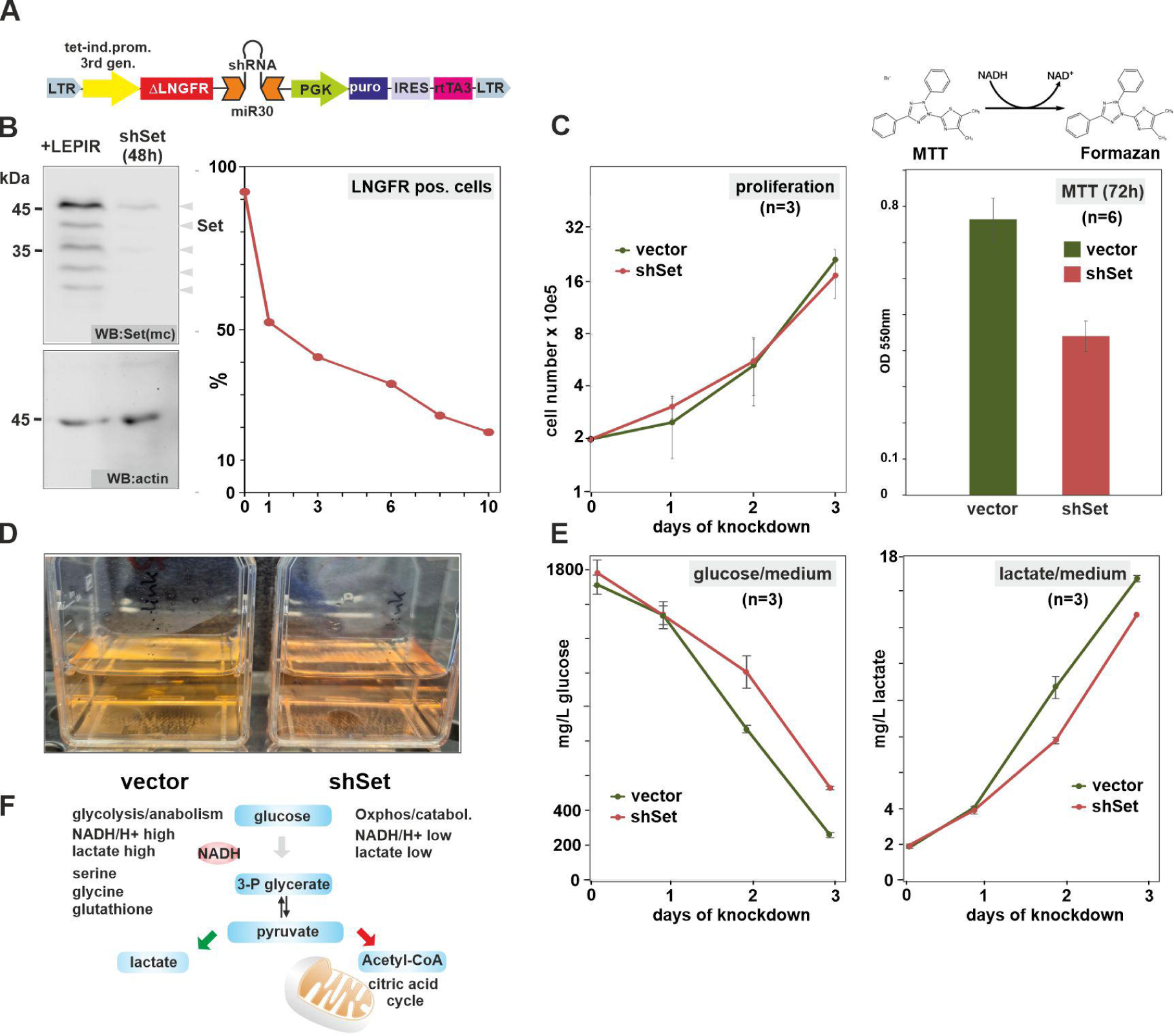
Set knockdown causes a metabolic phenotype. A: Structure of the inducible shRNA lentiviral construct pLEPIR B: Cells transduced with either a luciferase targeting (pLEPIR) or Set specific (shSet) lentivirus were induced with doxycycline and magnetically enriched for LNGFR positive (= shRNA expressing) cells. Extracts were analyzed after 48h by western (left panel) and by FACS analysis for LNGFR/shRNA positive cells in a time course (right panel). C: Proliferation of control (vector = pLEPIR) and shSet knock-down cells tested by cell count (left panel) and a standard MTT test (right panel). D: Visual aspect of vector and shSet transduced cell cultures. Equal numbers of each cell type were seeded and incubated for 24h. Medium color demonstrates differential acidification. E: Glucose consumption and lactate production are reduced after Set knockdown. Glucose (left) and lactate (right) levels were recorded by standard biochemical assays for a 72h time course in cultures seeded with equal amounts of control (vector) or Set knockdown (shSet) cells. F: Overview of the glycolysis/oxidative phosphorylation switch. Pyruvate kinase is a major regulator that allows cells to either emphasize glycolysis and lactate production for anabolic needs or to channel glycolytic products towards mitochondria for energy production.

### Set activates a pro-proliferative gene expression program

To gain insight into the global gene expression influenced by Set, we determined genome-wide transcription rates by nascent-RNA sequencing (also called SLAM-seq = thiol (SH)-linked alkylation for the metabolic sequencing). Vector control and Set knockdown cells were analyzed 48h after induction of shRNA expression. In total, 3217 genes were significantly downregulated after reduction of Set and 667 genes showed increased expression (threshold: log2_(Set on/Set off)_ >1 or <-1) (figure 4A, supplemental table 1). Although these genes likely also include secondary targets owing to the necessary time for a shRNA based knockdown to take effect, this points to a predominant role of Set as activator of transcription. Corroborating the metabolic phenotype observed before, both, *Hnrnpa1* and *Ptbp1* were sensitive to Set levels. Major Set-dependent genes coded for the ribosomal precursor transcript *Rn45s* and *Myc*, both hallmarks of proliferating cells. Transcripts upregulated in response to Set ablation belonged to glycolytic enzymes (Eno1, Aldoa, Ldha, Gapdh, Pkm) and multiple ribosomal subunits. We interpreted this as compensatory response of Set depleted cells to counter the suppression of glycolysis and ribosomal synthesis. Examination of clustered *HoxA* gene expression as sentinels of Mll involvement was hampered by the retroviral delivery of *HoxA9* for immortalization placing *HoxA9* under constitutive LTR control. Yet, the endogenous *HoxA1* gene and the divergently transcribed *Hotairm1* non-coding RNA negatively regulating *HoxA1* expression, did show a behavior consistent with Mll activity. While Set knockdown reduced *HoxA1*, *Hotairm* was induced (figure 4B). An unbiased analysis by GSEA (gene set enrichment analysis) confirmed our findings (figure 4C) with “ribosomal constituents” and “oxidative phosphorylation” gene sets strongly enriched in Set depleted cells. Conversely, the “constituent of chromatin” similarity, mainly driven by histone genes, argues for a role of Set in preparing cells for rapid proliferation.

**Figure 4:**
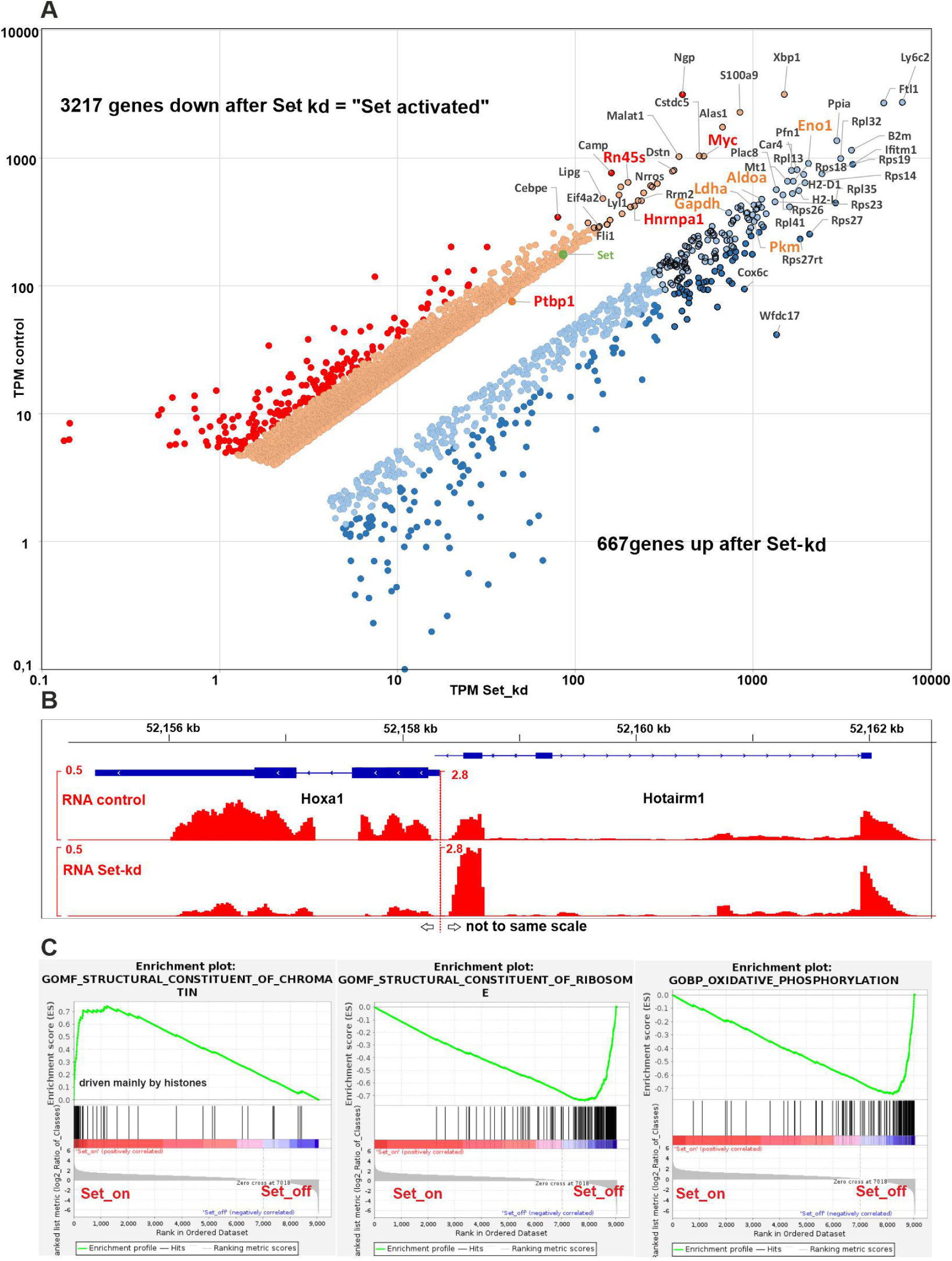
Set controlled gene expression program. A: Scatter plot depicting all transcripts with a significant change in transcription rate [log2 (Set_on_/Set_off_) > 1 or < −1] 48h after Set knockdown. Values were determined by nascentRNA sequencing and are given as TPM. The top 100 highest expressed transcripts are name-tagged. Key genes are emphasized in red. *Ptbp1* was plotted manually as it narrowly missed the threshold (see text). B: IGV snapshot of *HoxA1*/*Hotairm* transcripts in response to Set knock-down. C: Gene set enrichment analysis of the gene expression pattern responding to Set. The three top-scoring GSEA plots are shown.

### Set depletion impairs Mll binding and transcriptional elongation

To explore the function of Set in transcriptional regulation in detail, we performed ChIP for Mll, H3K4me3, total RNA PolII as well as the serine 2 and serine 5 phosphorylated versions of RNAPolII in vector control and shSet cells (figure 5A, B). Recording the changes observed for Set-controlled genes revealed that a reduction of Set induced a concomitant loss of Mll without affecting H3K4me3 modification. This indicated that Mll is not the main H3K4 methylase as reported before ^14^ and that H3K4 methylation is not necessarily proportional to final transcriptional output. In a similar direction, total RNAPolII and RNAPolII phosphorylated at serine 5 of the CTD (C-terminal domain, initiating polymerase) were almost unaffected by a Set knockdown while elongating RNAPolII Ser-2 P was selectively reduced. This aligns well with a proposed role of Mll predominantly in elongation.

**Figure 5:**
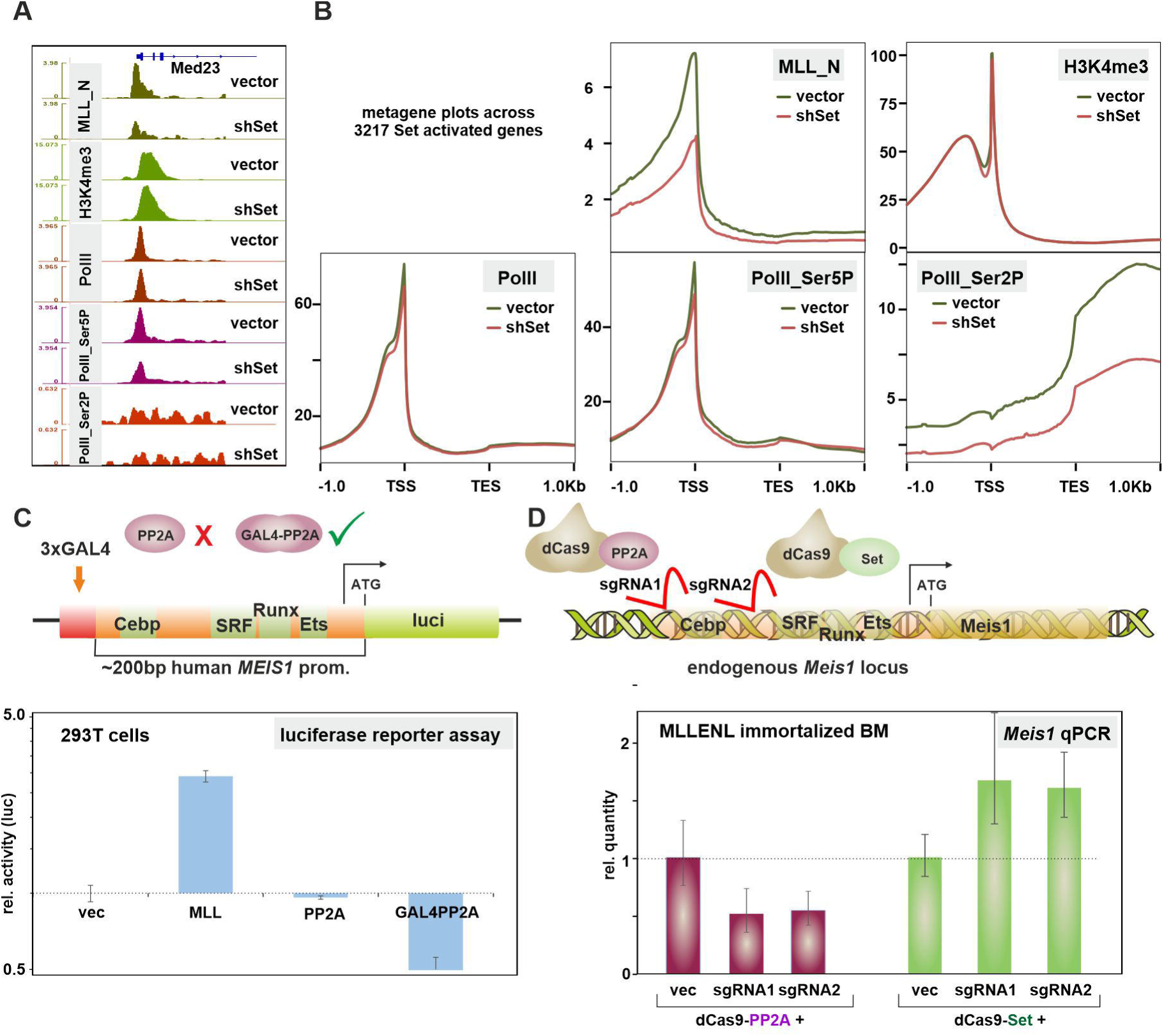
Set is required for Mll binding and elongation stimulation, while PP2A acts as repressor. A: IGV snapshot depicting ChIP profiles of Mll-N, H3K4me3, RNAPolII (total), RNAPolII-Ser5P, and RNAPolII-Ser2P at the promoter of the Set target *Med23*. B: Metagene plots across all Set activated genes reveal a specific loss of Mll and elongating RNA polymerase from chromatin after Set depletion. C: Only promoter recruited PP2A can repress the MLL responsive human Meis1 promoter. The reporter construct was co-transfected with expression constructs as indicated and luciferase results were determined in triplicate experiments. D: Recruitment of Set and PP2A to the endogenous *Meis1* promoter in myeloid precursor cells. Effectors were targeted to the murine *Meis1* promoter in MLL-ENL transformed cells by a fusion with catalytically inactive Cas9 (dCas9) in combination either with a non-targeting (vec) or two *Meis1* promoter specific sgRNAs. *Meis1* output was measured by RT-qPCR in triplicates.

Recently, nuclear PP2A accompanying active RNAPolII as member of the integrator complex has been described as negative regulator of transcriptional elongation ^15^. If PP2A localized at promoter regions together with Set/Mll would similarly affect transcription is unknown. To test this, we first resorted to an ectopic reporter system. The 200bp human MEIS1 core promoter ^16^ was cloned immediately downstream of a tandem triplicate of GAL4 DNA binding sequences in a standard luciferase reporter plasmid (figure 5C). First, we confirmed that this construct was responsive to co-expressed full length MLL in 293T cells. These cells express endogenous MEIS1 ensuring that all additional cellular factors necessary for MEIS1 promoter function are present. Overexpression of the catalytic subunit of PP2A alone did not alter reporter output confirming that dephosphorylation of cytoplasmatic or other nuclear proteins does not interfere with reporter expression. In contrast, recruiting PP2A to the promoter as GAL4 fusion protein caused a pronounced drop in reporter activity. Further support for a function of PP2A and Set in a natural promoter environment was obtained by introducing fusions of catalytically dead Cas9 enzyme (dCas9) with PP2A or Set into MLL-ENL transformed murine primary cells where, in contrast to HoxA9 transduced cells, endogenous *Meis1* is expressed. Cas9-PP2A and Cas9-Set were subsequently targeted to the endogenous *Meis1* promoter by two different sgRNAs or a non-targeting decoy followed by *Meis1* RT-qPCR (figure 5D). Although effect sizes were moderate, clear transcriptional effects were recorded supporting PP2A as inhibitor and Set as activator, respectively.

### Set is necessary to recruit active Msk1 to promoters

In a search for molecules that might be potential targets protected by Set from dephosphorylation we concentrated on Msk1 (mitogen and stress activated kinase 1) because Msk1 has been associated before with MLL and MLL fusion proteins ^17^ and it is phosphorylated by Erk1/2 for activation ^18^. ChIP with a phospho-specific antibody for Msk1 Ser376P was performed in control and Set knockdown cells (figure 6A, B, C). Msk1-P distribution did not only perfectly colocalize with MLL and Set, but presence of Msk1-P was also dramatically reduced after reduction of Set concentrations. Interestingly, H3S10 phosphorylation, a known catalytic target of Msk-1, did neither respond to Set nor was it generally detectable around promoter regions. This suggested a different function for Msk1 in promotor regions. As tethering MLL to chromatin requires menin we checked the interaction of Msk1 with MLL and menin by co-immunoprecipitations (figure 6E, F). These experiments clearly indicated a direct interaction of Msk1 with MLL as well as with menin. In summary, these results are best explained by the presence of a MLL/menin/Msk1 complex at promoter regions whose assembly is regulated by phosphorylation.

**Figure 6:**
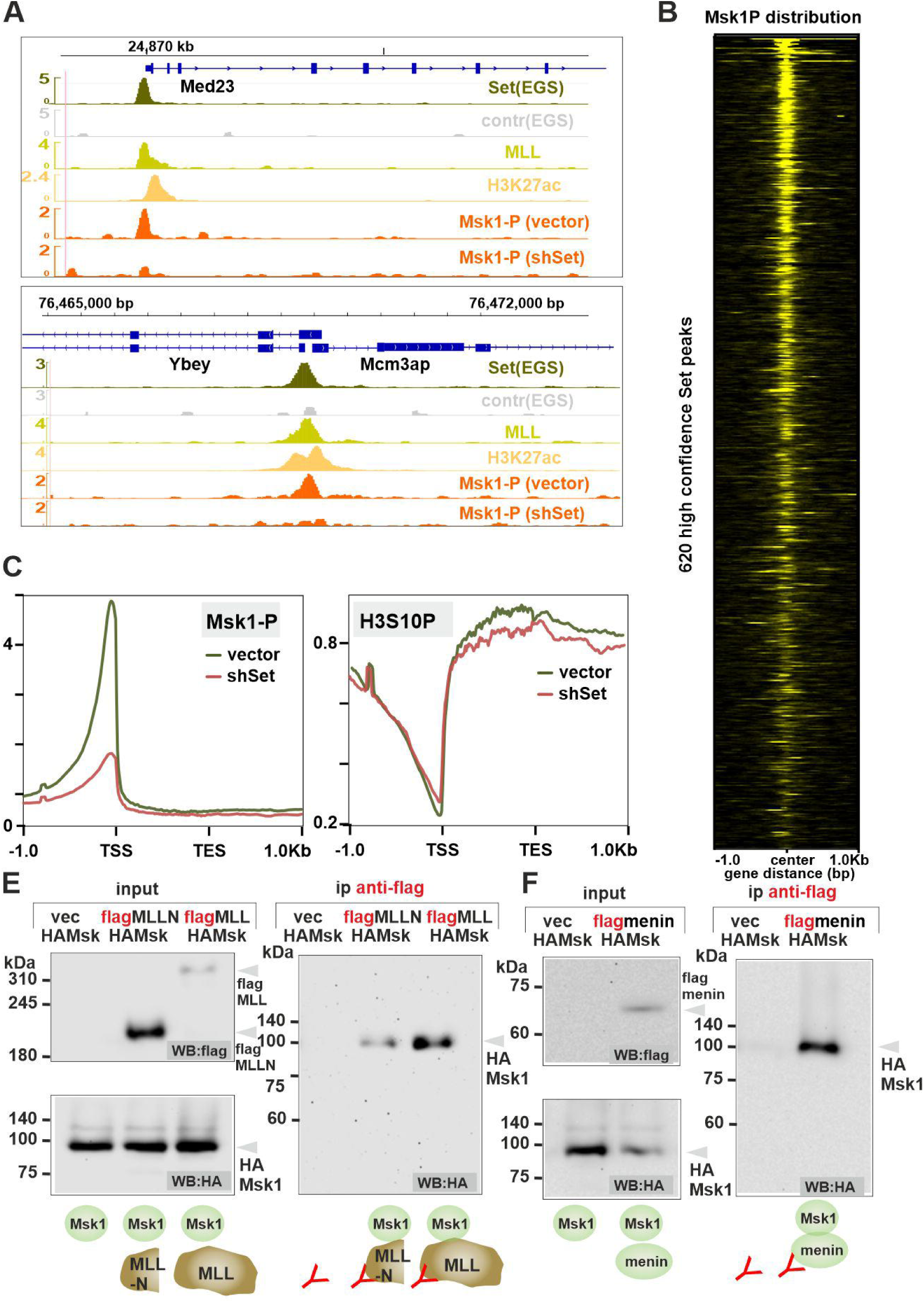
Msk1 forms Set dependent complexes with Mll and menin at promoters. A: IGV snapshot of the distribution of active Msk1 (serine 133 phosphorylated) on H3K27acetylated promoter regions co-occupied by Mll on Set targets *Med23* and *Ybey*. B: Heat map of Msk1-P binding around identified Set peaks. C: Metagene plot of Msk1-P and phospho H3S10, a known Msk-1 catalytic product around Set regulated genes in control and Set knockdown cells. D: Direct interaction of Msk1 and Mll/Set as evidenced by co-immunoprecipitations of tagged versions expressed in 293T cells.

### Set is required to establish a promoter specific acetylation pattern

While mechanistic details of elongation stimulation by MLL are largely unknown, we surmised that histone acetylation must be important in this process. For one MLL associates with histone acetyltransferases ^19–21^ and second, transcriptional elongation complexes like P-TEFb (positive transcription elongation factor b) are recruited by YEATS-domain containing proteins. YEATS domains, in turn, are readers of histone acetylation ^22^. To further investigate the process that relays a loss of Set/Mll to a reduction of RNAPolymerase Ser2-phosphorylation, we performed ChIP for H3K9ac, H3K14ac, and H3K27ac commonly found in promoter regions (figure 7A, B upper panel). A knockdown of Set and therefore a loss of Mll almost exclusively affected H3K14ac with only marginal reduction in H3K9ac and H3K27ac revealing an unexpected specificity. In total up to 14 different histone acetyltransferases (HAT) are known to acetylate H3K14 ^23^. To begin to explore the responsible HATs we extended ChIP to Cbp (and its binding partner Creb in the active, serine 133 phosphorylated version) as well as Moz, both of which have been linked to MLL (figure 7B lower panel). Set knockdown did not affect Cbp and Creb-P was even increased after Set depletion, suggesting a possible compensatory effect. In contrast, Moz occupancy paralleled Set reduction, however to a smaller extent than H3K14ac itself, implying further HAT activities involved in setting this mark.

**Figure 7:**
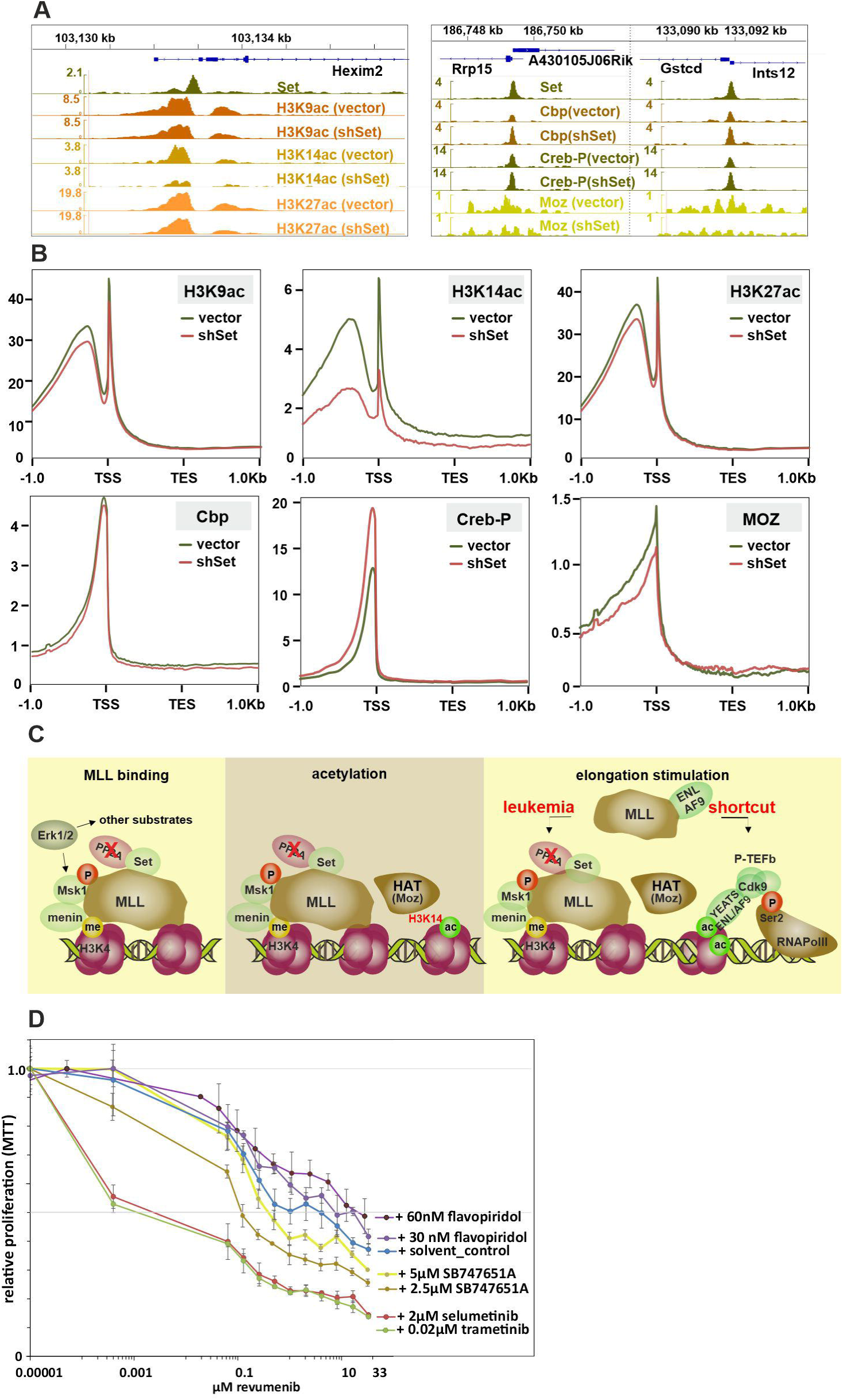
Mll induced H3K14 acetylation requires Set; cooperation of signaling kinase and menin inhibitors. A: IGV snapshots of acetylation pattern as well as binding of HATs Cbp/Creb-P and Moz at promoter regions of select Set targets. B: Metagene plots of the same distributions across all Set regulated genes. C: Schematic depiction of Mll as signal integrator. Signaling establishes a proper phosphorylated environment protected by the anti-phosphatase Set as prerequiste for Mll localization to promoters. Once bound, Mll establishes H3K14acetylation likely involving Moz. Acetylation can be read by YEATS domain proteins recruiting RNAPolII Ser2 kinase containing complexes like PTEF-b. While Mll fusions shortcut the need for HAT activity by directly recruiting elongation activities, the need for signaling to permit chromatin localization is conserved. D: Cooperation of various kinase inhibitors with the menin inhibitor revumenib. The IC50 of revumenib was determined in 96h standard MTT assays in the presence of kinase inhibitors supplied at approximately their individual IC50. Flavopiridol = Cdk9 inhibitor, SB747651A = Msk1 inhibitor, Selumetinib/Trametinib = Erk1/2 inhibitors.

### Kinase and menin inhibitors cooperate to block activity of a MLL fusion protein

A loss of Set or an activation of PP2A by FTY720 has been previously shown to impair MLL-fusions ^11, 12^. MLL joined to a member of elongation complexes would effectively shortcut the requirement to establish a specific acetylation pattern (for a schematic depiction see figure 7C) explaining the runaway elongation seen in MLL-r cells ^24^. The Set/phosphorylation dependent targeting of MLL/menin/Msk1 to chromatin, however, is a functional dependency that should be retained for MLL fusions. Therefore, we tested, if inhibitors of Msk1 (SB747651A) or inhibitors of Erk1/2 as activators of Msk1 (selumetinib, trametinib) would cooperate with menin inhibitors to block MLL-ENL induced transformation. A Cdk9 inhibitor (flavopiridol) was used as control. Two inhibitors that act on the same pathway with one epistatic towards the other will show additive behavior, whereas inhibitors that target independent activities can be expected to act cooperatively. Therefore, we determined IC50 concentrations for the menin inhibitor revumenib in the presence of other kinase inhibitors in MLL-ENL transformed primary murine cells (figure 7D). Cooperativity would be indicated by a reduction of IC50 concentrations for menin inhibition, whereas additive activity should not affect this value. Flavopiridol, blocking Cdk9 downstream of MLL-ENL was used as proof of principle. Corroborating our assumptions, flavopiridol did not significantly alter revumenib IC50 concentrations. Conversely, SB747651A, a Msk1 inhibitor, particularly at low levels reduced revumenib IC50 values slightly, indicating some level of cooperativity. Remarkably, however, ablating Erk1/2 signaling with selumetinib or trametinib strongly cooperated with menin inhibition. In summary, these results indicate that also MLL fusions rely on signaling kinase input for their function.

## Discussion

Here we pinpoint phosphorylation and its protection from phosphatase erasure by Set as an important feature that is prerequisite for binding of a Mll/Msk1/menin (MMM) complex to promoter chromatin. Once bound, MMM establishes a specific acetylation pattern characterized by H3K14ac that finally translates into recruitment of RNAPolII Ser2-kinase containing elongation complexes. Some molecular details, however, are still missing. We do not know the actual phosphorylation targets beyond Msk1 itself. According to high throughput proteomic databases (phosphosite.org), MLL and menin are phosphorylated at multiple residues. It will require future research to determine if these proteins indeed are substrates of Msk1 or other kinases and if modification is necessary for binding of MMM to chromatin. Moreover, additional, less researched proteins involved in MLL localization like LEDGF ^25^ may play a role.

Yet, even without these details, our results emphasize an important and hitherto largely neglected aspect of MLL biology. MLL is a relay station for cellular signaling. While it is self-evident that the transcriptional regulators necessary for expression must be present *before* the respective gene is activated, it has received little mechanistic attention how all these factors are kept inactive while not actively engaged. An obvious solution is activation by kinase signaling. Our results characterize Set as a signaling “enhancer” protecting the necessary phosphorylation required to tether MMM to chromatin. This goes well along with observations that overexpression of SET either by genetic alterations or by a block of SET-degradation through SETBP1 is leukemogenic ^1^. The major MLL targets, the HOX homeobox proteins are key regulators of normal and malignant hematopoiesis ^26, 27^. HOXA9 is the single most prognostic factor in leukemia associated with dismal survival. A role for SET/signaling in MLL function fits well with the observation that myelodysplastic and myeloproliferative disorders frequently not only harbor mutations in signal molecules but they also feature alterations of SETBP1. In a similar direction, experimental overexpression of *Setbp1* is accompanied by massive upregulation of *HoxA9* and *HoxA10* ^9^. Furthermore, NUP fusions localize to *HOX* loci ^28^ providing an explanation how NUP214-SET fusions stimulate *HOX* transcription through MLL and cause leukemia. As an interesting side aspect, *SETBP1* mutations are also detected in neurological disease ^29^ paralleling the fact that altered transcriptional elongation frequently manifests itself in neurological abnormalities as exemplified by mutations of *Mllt1*/*Af4* that cause the “robotic” phenotype in mice ^30^. Although MLL fusion proteins short-circuit the need for acetylation to induce transcriptional elongation they still seem subject to phospho-dependent location control. That makes signaling kinase inhibitors as complement or even replacement for menin inhibitors a promising perspective for further research.

## Methods

### DNA, cells, inhibitors, antibodies, and degron constructs

Proteins were expressed from cloned cDNAs inserted either into pMSCV retroviral backbones (Clontech, Palo Alto, CA) or in pcDNA3 based expression vectors (ThermoFisher, Germany). All insert sequences were derived from laboratory stocks or assembled from cDNA by PCR with confirmation by sequencing. In our hands, clones containing *Set* cDNA sequences were very prone to recombination in *E.coli* strains. Therefore, for all cloning procedures involving *Set* we used the CopyCutter strain (Avantor Europe, Gliwice, Poland) that suppresses copy numbers of plasmids. PROTAC (proteolysis targeted chimaera) modified proteins were obtained by fusion with a F36V mutated FKBP degron. Rapid degradation is induced by addition of the heterobifunctional dimerizer dTAG13. This forces dimerization with the endogenous E3-ligase cereblon tagging the FKBP^F36V^ fusion protein for destruction. As demonstrated previously by mass spectrometry ^31^, dTAG13 has no discernible effect on endogenous proteins. For simplicity FKBP^F36V^ will be denoted FKBP throughout the text.

Primary cells used in this study were derived from CD117 (Kit) positive HSPCs isolated from bone marrow of C57/BL6 mice with a triple-ko for granule protease genes *Elane*, *Prtn3*, and *Ctsg* ^32^. The protease deficient environment does not cause hematopoietic abnormalities but avoids protein degradation during the ChIP procedure in myeloid precursor cells. HSPCs were immortalized by transduction with HoxA9 or MLL-ENL or the degradable derivative HoxA9-FKBP. Inducible knockdown of endogenous Set was achieved by super-transducing cells with a lentiviral construct (pLEPIR) targeting the sequence AACAGAATTTGAAGACATTAAA in the Set cDNA. pLEPIR is a laboratory modification of the original mir-E construct described by Fellman et al ^33^. For initial establishment cells were cultivated in methylcellulose (M3534, StemCellTechnologies, Cologne, Germany) for two rounds under antibiotics selection. Subsequently cultures were explanted and maintained in RPMI1640 (Thermo-Scientifc, Germany) supplemented with 10% FCS, penicillin-streptomycin, 5ng/ml recombinant murine IL-3, IL-6, GM-CSF, and 50ng/ml recombinant murine SCF (Miltenyi, Bergisch-Gladbach, Germany). dTAG13 was from Tocris (NobleParkNorth, Australia). All other chemicals were provided either by Sigma (Taufkirchen, Germany) or Roth (Karlsruhe, Germany).

### ChIP-Seq, cell lysis, nascent-RNA isolation

ChIP was performed as described in ^34^ using material crosslinked for 10 min in 1% formaldehyde at RT. For ChIP of Set and Moz a modified crosslinking protocol was used. Washed cells were incubated for 30min at room temperature in 2mM EGS (Ethylenglycol bis (Succinimidyl-Succinat, Thermo-Fisher, Germany) followed by a 10min additional crosslink in 1% formaldehyde. ChIP lysis was done in 50mM Tris/HCl pH8.0, 10mM EDTA, 100mM NaCl, 1mM EGTA, 0.1% sodium-deoxycholate, 0.5% N-lauroylsarcosine, 1mM PMSF and 1% HALT complete protease/phosphatase inhibitor cocktail (Pierce, Thermo-Fisher, Germany). Precipitation was performed from 5×10e6 cells per experiment using protein G coupled paramagnetic beads (Cell Signaling Technologies). Antibodies used for ChIP and western blots were purchased from Cell Signaling Technologies (CST) and used at 5µg per assay if not stated otherwise. Set : Invitrogen (#MA5-35772); MLL: CST (#8178) mixed 1:1 with anti-MLLN, Upstate (Temecula, CA, #05-764) 10 µl AB-mix per 5×10e6 cells, H3K4me3: CST (#9751), RNAPolymeraseII: CST (#14958), RNAPolII-Ser5P: CST (#13499), RNAPolII-Ser2P: CST (#13523), Msk1-P (Ser376P): CST (#9591), H3S10P: CST (#53348), H3K9ac: CST (#9649), H3K14ac: CST (#7627), H3K27ac: CST (#8173), Cbp: CST (#7389), Creb-P (Ser133P): CST (#9198), MOZ: Invitrogen (PA5-68046) mixed 1:1 with Invitrogen (PA5-103467), HA: CST (#3724), flag: Merck (F1804).

Cell lysis for western was done in 20mM HEPES pH 7.5, 10mM KCl, 0.5mM EDTA, 0.1% triton-X100 and 10% glycerol supplemented with 1mM PMSF and 1% HALT complete protease inhibitor (triton lysis). Nascent-RNA isolation was done exactly as described in ^6^. Coimmunoprecipitations were performed with total lysates of transfected 293T cells extracted with triton buffer as above supplemented with 300mM NaCl. Material bound to protein-G beads was washed extensively with triton-lysis buffer containing 150mM NaCl before western analysis.

### NGS and bioinformatics

Sequencing libraries were either prepared using NEBNext Ultra™ II DNA Library Prep Kit reagents (NEB, Ipswitch, MA) or with NEBNext Single Cell/Low Input RNA Library Prep reagents according to the manufacturer. Size selection for ChIP libraries was done on final libraries. Sequencing was done at the in house core facility as 150bp paired end reads according to standard Illumina pipelines.

Data were mapped with BWA mem (0.7.17) ^35^ to the *Mus musculus* mm10 genome. Reads mapping more than once were excluded by filtering for sequences with a mapping quality score > 4. For visualization BAM files were normalized and converted to TDF format with IGV-tools of the IGV browser package ^36^. Peak finding, motif analysis and peak annotation was done with Homer (4.9.1) ^37^. BAM files were converted to bigwig by Deeptools (3.0.0, bamCoverage) ^38^. Metagene plots were created with Deeptools (3.0.0). Matrices were calculated with calculateMatrix and plotted with plotHeatmap from the Deeptools suite. RNA derived reads were aligned with STAR (v020201) ^39^ to the reference genome mm10 and reads derived from repetitive sequences were excluded by samtools (view)1.8 ^40^. Transcripts were quantified by Homer (v5.0) analyzeRepeats routines and further evaluated with standard spreadsheet tools.

### Statistics

Where appropriate two-tailed T-test statistics were applied.

## Data sharing

NGS reads are available with the European Nucleotide Archive under accession number PRJEB107176

## Acknowledgements

We thank Renate Zimmermann for technical assistance and Prabha Varghese for help with cloning. This work was supported by research funding from Deutsche Forschungsgemeinschaft grant SL27/9-2 and in part by Deutsche Krebshilfe grant 70114166 both awarded to RKS. All authors declare no conflict of interest

## Author contributions

MPGC, and RKS performed and analyzed experiments. RKS performed NGS data analysis, conceived and supervised experiments, RKS wrote the manuscript. All authors read and discussed the manuscript.

